# Multi-site phosphorylation of the TOPBP1 ATR Activation Domain propels ATR signaling during the response to DNA breaks

**DOI:** 10.1101/2022.09.19.508511

**Authors:** Holly Senebandith, Kenna Ruis, Katrina Montales, Oanh Huynh, Yasaman Jami-Alahmadi, James A. Wohlschlegel, W. Matthew Michael

## Abstract

TOPBP1 is a BRCT domain-containing scaffold protein that plays important roles in a diverse array of cellular processes, including DNA damage signaling, DNA repair, DNA replication, and mRNA transcription. For DNA damage signaling, TOPBP1 activates the crucial damage response kinase ATR and this occurs during replication stress as well as during a DNA double-strand break response. ATR signaling allows cells to survive genotoxic issues and represents a formidable barrier to transformation to a tumorigenic state. Despite its importance to genome stability, the biochemical mechanism for how TOPBP1 activates ATR is not fully understood. TOPBP1 uses a discrete domain, termed the ATR activation domain, to stimulate ATR kinase. Recent work has shown that the AAD must be in a multimeric state to activate ATR. Other work has shown that phosphorylation of the AAD on serine 1131 (in Xenopus) is important for its function, and some have suggested that this is linked to the formation of TOPBP1 condensates during a DNA damage response. In this study we examine AAD phosphorylation in detail and we report three important new findings. One, S1131 phosphorylation promotes ATR activation in a manner independent of condensate formation and is instead linked to promoting multimerization of the AAD. Two, we identify a novel sight of AAD phosphorylation, on T1098, and show that is required for ATR activation. Three, we identify additional, candidate phosphorylation sites, some of which fit the consensus for casein kinase 2, and we show that casein kinase 2 activity is required for the AAD to perform its function. These studies show that multi-site phosphorylation of the AAD is an important component of the mechanism by which TOPBP1 activates ATR.

## 1. Introduction

Ataxia telangiectasia and Rad3-related (ATR) kinase is a vital component of the cellular response to replication stress and DNA damage (Salvidar et al, 2017; Waterman et al, 2020). ATR is also important for the proper timing of cell cycle progression under unperturbed conditions (Saldivar et al, 2018). This large phosphatidylinositol 3-kinase-like kinase (PIKK) is hyper-activated at sites of DNA damage and it goes on to phosphorylate hundreds of substrates to allow cells to survive genotoxic stress. ATR signaling is also a barrier to cancer as it will drive damaged cells into senescence, thereby preventing their continued proliferation (Bartkova et al. 2006; Di Micco et al. 2006). Despite the importance of ATR signaling to human health and disease, it remains unclear how ATR is activated by damaged DNA or stalled replication forks.

So far, two proteins have been identified that help activate ATR at sites of damage, TOPBP1 and ETAA1 (reviewed in Niedzwiedz, 2016; Salvidar et al, 2017). The work presented here focuses on TOPBP1, which is composed of nine BRCT domains and a discrete and stand-alone ATR activation domain (AAD; Figure 1A). To initiate DNA damage signaling, for example at sites of DNA double-strand breaks (DSBs), ATR is recruited to the DSB via its binding partner ATRIP while TOPBP1 is recruited separately, via an unknown mechanism. At the DSB, TOPBP1 undergoes a conformational change that allows its AAD to contact ATR and ATRIP, in a manner that greatly stimulates ATR kinase activity. Recent work from our group, and others, has revealed that the conformational change is linked to multimerization of TOPBP1 at sites of damage (Kim et al, 2020; Thada et al, 2021; Ruis et al, 2021). Notably, while the AAD by itself can activate ATR when present at high concentrations, forced multimerization of the AAD renders it a far more potent activator (Kim et al, 2020; Thada et al, 2021). Indeed, forced multimerization of the full-length TOPBP1 protein also produces more robust ATR activation (Zhou et al, 2013). These studies make it clear that the way TOPBP1 is regulated is through control of its oligomeric state; TOPBP1 only assumes a multimeric state when it is bound to damaged DNA, and thus ATR is only activated at sites of damage and is not constitutively activated in solution.

**Figure 1.**
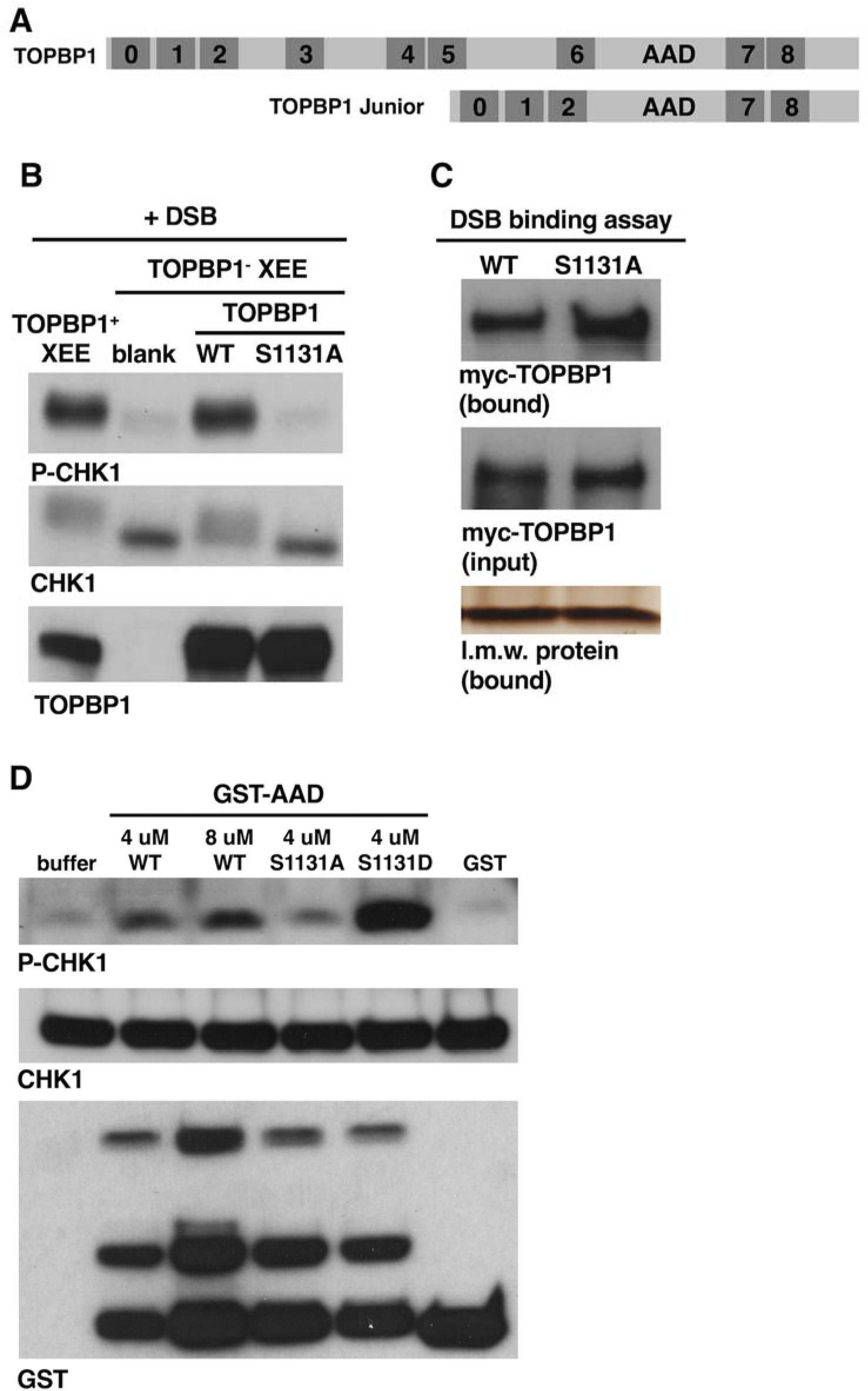
S1131 in TOPBP1’s AAD is dispensable for recruitment of TOPBP1 to DSBs but is required for ATR activation in the DMAX system. **A.** Schematic representation of full length TOPBP1 and TOPBP1 Junior. **B.** A DMAX assay where XEE was immunodepleted of endogenous TOPBP1 and then supplemented with the indicated IVTT-produced TOPBP1 proteins prior to the addition of DSBs. TOPBP1^+^ XEE (lane 1) refers to extract that was mock-depleted using non-specific IgG. Blank (lane 2) refers to a TOPBP1-depleted extract that received an unprogrammed IVTT reaction. After incubation, the samples were probed by Western blotting for P-CHK1, CHK1, and TOPBP1 (using anti-TOPBP1 antibody HU142). The experiment shown is representative of two independently performed biological replicates. **C.** HSS was combined with IVTT-produced and myc-tagged TOPBP1 proteins. These proteins were the wild type form (“WT”) and the S1131A mutant. DSB beads were then added and the samples were incubated for 30 min. After incubation, the beads were washed and eluted and the samples were probed by Western blot for the indicated protein. The myc antibody was used to probe for TOPBP1. The bound samples were also stained by silver and a 12.5 kDa low molecular weight (l.m.w) protein is shown, to demonstrate equal isolation of the beads. The experiment shown is representative of two independently performed biological replicates. **D.** The indicated GST proteins were added to XEE at the indicated concentrations and incubation was carried out for 30 minutes at room temperature. The samples were then probed for P-CHK1, CHK1, and GST. The experiment shown is representative of two independently performed biological replicates.

These findings have now focused attention on how TOPBP1 multimers form at sites of damage. Recent work from our group has shed light on this with the derivation of a synthetic form of TOPBP1 that is a stripped-down version lacking all sequences irrelevant to ATR signaling (Ruis et al, 2021). This protein, termed TOPBP1 Jr, contains (1) the first three BRCT domains, which are needed for recruitment to DSBs, (2) the AAD, and (3) the final two BRCT domains, which are required for multimerization (Figure 1A). Other works have shown that sequences within the AAD itself are important for regulation of TOPBP1. For example, it has been shown that serine 1131 in Xenopus TOPBP1 must be phosphorylated by the ataxia telangiectasia mutated (ATM) kinase in order for TOPBP1 to activate ATR (Yoo et al, 2007). Other, more recent work has suggested that phosphorylation of this same serine residue in human TOPBP1 causes the protein to undergo liquid-liquid phase separation to create TOPBP1 condensates at sites of damage (Frattini et al, 2021). These condensates are micron-sized, contain millions of TOPBP1 molecules, and are proposed to amplify ATR signaling.

In this work we use our <u>D</u>SB-<u>m</u>ediated <u>A</u>TR activation in *<u>X</u>enopus* egg extract system (DMAX; Montales et al, 2021; Ruis et al, 2022; Montales et al, 2022) to study how TOPBP1’s AAD is regulated. DMAX offers several powerful advantages for mechanistic studies of ATR signaling. For example, ATR signaling is readily assessed by scoring for phosphorylation of the key ATR substrate CHK1. Recruitment to DSBs is also directly assayed in DMAX using DSBs that are immobilized on magnetic beads in a pull-down assay. Structure-function analysis is another key feature of DMAX as target proteins can be removed from *Xenopus* egg extract (XEE) via immunodepletion and mutant forms, rapidly and easily produced by *in vitro* transcription and translation (IVTT), are added back for phenotypic assessment. Using DMAX, we show here that S1131 in the AAD plays an important role in TOPBP1 multimerization independently of condensate formation. We also identify a novel site of phosphorylation within the AAD, T1098, and show that this residue is required for ATR activation. Lastly, we provide evidence that casein kinase 2 is required for ATR activation.

## 2. Material and Methods

### 2.1 Materials

#### 2.1.1 Plasmids

All plasmids used in this study are described below, in table format, and all are *Xenopus* proteins. Specific details about plasmid construction are available upon request.

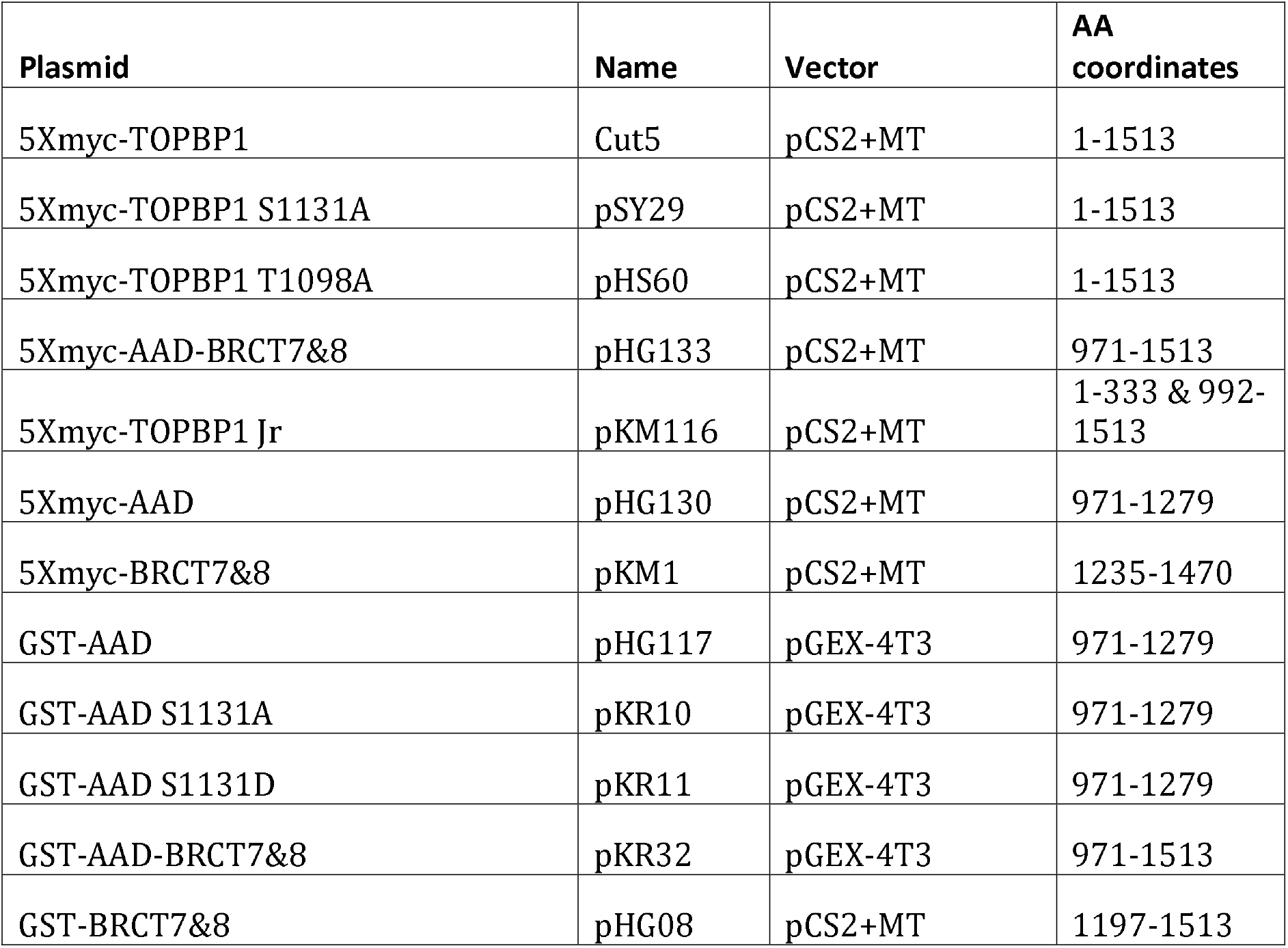

#### 2.1.2 Recombinant proteins

The recombinant proteins used in this study were GST, GST-AAD (wild type, S1131A, S1131D), GST-BRCT7&8, GST-AAD-0BRCT7&8). All proteins were expressed in *E.coli* BL21(DE3) cells, at 37°C for 4 hours, and all proteins were purified from the soluble fraction according to standard procedures (detailed below).

#### 2.1.3 Antibodies

We used the following commercially sourced antibodies in this work: Myc (Millipore Sigma #M4439) and GST (Millipore Sigma #05-782). We also used our own antibody against *Xenopus* TOPBP1, HU142, which has been described (Van Hatten et al., 2002).

### 2.2 Methods

#### 2.2.1 Xenopus egg extracts

The high-speed supernatant (HSS) of XEE was used exclusively in this study. HSS was prepared exactly as described (Smythe and Newport, 1991). For immunodepletion of TOPBP1, the HU142 antibody was used and the procedure was performed exactly as described (Van Hatten et al, 2002). Depleted XEEs were then supplemented with IVTT-produced proteins (2.5 μl IVTT lysate in 20 ul of XEE) as described (Montales et al, 2021).

#### 2.2.2 IVTT production of proteins and determination of protein concentrations

IVTT reactions were performed using the SP6 TnT® Quick Coupled Transcription/Translation System (Promega #L2080) according to the manufacturer’s instructions.

#### 2.2.3 Recombinant Protein Expression and Purification

Plasmids were transformed in BL21 (DE3) Competent *E. Coli* cells and grown in LB media (GST fusion proteins) or LB media with 0.2% glucose (MBP fusion proteins) with ampicillin. At an OD600 of 0.6-0.8, cells were induced with 0.3 mM IPTG for 4 hours at 37 C. Cells were then harvested by centrifugation (4000 rpm for 20 min) and washed with phosphate-buffered saline (PBS). Cells were lysed by sonication in GST column buffer (40% PBS, 60% TEN buffer: 10 mM Tris-Cl, pH 8.0, 1 mM EDTA, pH 8.0, 100 mM NaCl) supplemented with cOmplete protease inhibitor cocktail tablets (Roche #04693159001) and 10 mg lysozyme. Lysates were cleared by centrifugation (15,000 rcf for 20 min) and incubated with glutathione sepharose (BioVision #6566) for 1 hour at 4 C. Beads were poured over a column and washed with column buffer. GST fusion proteins were eluted with 20 mM glutathione in 75 mM HEPES, pH 7.5, 150 mM NaCl, 5 mM DTT, and 0.1% Triton-X. Proteins were dialyzed overnight into PBS.

#### 2.2.4 ATR activation (DMAX) assays

For DMAX assays, okadaic acid was first mixed with 20 μl of HSS to a final concentration of 1 μM, as described (Montales et al, 2021). Linear dsDNA derived from EcoRI-digested lambda DNA (Montales et al, 2021) was then added to the mixture and reactions were incubated at room temperature for 30 min. Samples were analyzed via Western blotting using standard conditions. In some cases, when GST-AAD proteins were used, then no DNA was included in the assay. For the experiments shown in Figure 4, the casein kinase 2 inhibitor CX-4945 was used, and it was purchased from https://Selleckchem.com (catalog # S2248).

#### 2.2.5 DSB binding assay

These were performed exactly as described (Montales et al, 2021). Briefly, a 5 kb PCR fragment was biotinylated via inclusion of a biotin moiety on the forward PCR primer. After PCR and cleanup, the 5 kb PCR fragment was coupled to magnetic streptavidin beads (Dyna-beads M-270 Streptavidin, ThermoFisher) according to the manufacturer’s instructions. These “DSB beads”, containing 600 fmol of dsDNA per assay, were then incubated in 20 μl of HSS supplemented with 5 μl of an IVTT reaction programmed to produce the protein of interest. After incubation the beads were collected on a magnetic stand and washed three times in PBS + 0.1% TritionX-100. Bound proteins were then eluted with 2X SDS-PAGE sample buffer and examined by Western blotting.

#### 2.2.6 Protein binding assays

For the GST pull-down assays, 5–10 ug of GST fusion proteins were bound to glutathione sepharose beads (BioVision #6566) in Binding Buffer (PBS + 0.1 % NP-40) for one hour at 4°C and then the beads were washed with Binding Buffer. Next, 20 μL of IVTT proteins were diluted into 400 μL of Binding Buffer and then incubated with the washed beads for one hour at 4°C. The beads were washed four times with 500 μL of Binding Buffer and then eluted with 2X Sample Buffer (Millipore Sigma #S3401-10VL). The input and bound fractions were then analyzed via Western blotting using standard conditions.

##### Mass spectrometry analysis of GST-AAD

GST-AAD was incubated in XEE for 30 minutes at room temperature and purified back out of the extract via glutathione-sepharose affinity chromatography. The sample was run on SDS-PAGE and a Coomassie-stained band was excised from the gel. Proteins in coomassie-stained gel slices were reduced, alkylated and digested by trypsin as previously described (Kaiser and Wohlschlegel, 2005). After digestion, peptides were extracted from the gel slices using 100 μl of 60% acetonitrile/0.1% TFA, dried in a SpeedVac vacuum concentrator, desalted using C18 tips, and reconstituted in 10 μl of 5% formic acid for LC-MS/MS analysis.

Desalted peptide mixtures were loaded onto a 25 cm long, 75 μm inner diameter fused-silica capillary, packed in-house with bulk 1.9 μM ReproSil-Pur beads with 120 Å pores as described previously (Jami-Alahmadi et al, 2021). Peptides were analyzed using a 140 min water-acetonitrile gradient delivered by a Dionex Ultimate 3000 UHPLC (Thermo Fisher Scientific) operated initially at 400 nL/min flow rate with 1% buffer B (acetonitrile solution with 3% DMSO and 0.1% formic acid) and 99% buffer A (water solution with 3% DMSO and 0.1% formic acid). Buffer B was increased to 6% over 5 min at which time the flow rate was reduced to 200 nl/min. A linear gradient from 6-28% B was applied to the column over the course of 123 min. The linear gradient of buffer B was then further increased to 28-35% for 8 min followed by a rapid ramp-up to 85% for column washing. Eluted peptides were ionized via a Nimbus electrospray ionization source (Phoenix S&T) by application of a distal voltage of 2.2 kV.

All mass spectrometry data were collected using data dependent acquisition on Orbitrap Fusion Lumos Tribrid mass spectrometer (Thermo Fisher Scientific) with an MS1 resolution of 120,000 followed by sequential MS2 scans at a resolution of 15,000. The Integrated Proteomics pipeline 2 was utilized to generate peptide and protein identifications. MS2 spectra were searched using the ProLuCID algorithm against the Xenopus reference proteome (xenbase.org) followed by filtering by DTASelect using a decoy database estimated false discovery rate of < 1% (Xu et al, 2015; Cociorva et al, 2007).

## 3. Results and Discussion

### 3.1 Serine 1131 promotes TOPBP1’s ability to activate ATR in a manner independent of condensate formation

Previous work from the Dunphy group had identified S1131 in TOPBP1’s AAD as an ATM phospho-acceptor site that is required for ATR activation in XEE under some conditions (Yoo et al, 2007). S1131 is required for ATR activation by small homopolymeric oligonucleotides like A_70_-T_70_ (AT70), and by sperm chromatin digested with EcoRI to produce DSBs. Interestingly, S1131 is dispensable for ATR activation by stalled replication forks. We thus initiated our studies by checking the requirement for S1131 in our DMAX system. To do so, we performed immunodepletion, where either non-specific or anti-TOPBP1 antibodies are coupled to Protein A beads and then incubated with XEE. After incubation, the XEE is recovered away from the antibody beads containing the captured TOPBP1 and used for ATR activation assays. EcoRI-digested λ DNA was used as a source of DSBs. We also prepared recombinant forms of TOPBP1, using IVTT in rabbit reticulocyte lysates, so that they could be added back to the depleted extracts. As shown in Figure 1B, a mock-depleted extract (treated with non-specific antibody beads, lane 1, labeled “TOPBP1^+^ XEE”) contained TOPBP1 and was able to activate ATR. By contrast, for the TOPBP1-depleted extract, TOPBP1 levels were substantially reduced, and this extract was deficient for ATR activation (Figure 1B, lane 2). When wild type TOPBP1 was added back to the TOPBP1-depleted extract, then ATR activation was restored (lane 3), and this did not occur when an S1131A mutant form of TOPBP1 was added back (lane 4). This experiment shows that S1131 is required for ATR activation by TOPBP1 in our DMAX system. Previous work had not addressed if S1131 is important for TOPBP1 recruitment to DSBs, and thus we employed our DSB binding assay to examine this important issue. DSB beads were prepared as previously described (Montales et al, 2021), where linear, 5kb dsDNAs with biotin moieties attached to end are immobilized on magnetic streptavidin beads. The beads are then incubated in XEE containing myc-tagged forms of either wild type or S1131A TOPBP1. Since our previous work has repeatedly shown that TOPBP1 does not bind to streptavidin beads lacking DNA under these assay conditions (Montales et al, 2021; Ruis et al, 2022; Montales et al, 2022), we did not include empty beads in this experiment. After incubation, the beads were isolated, washed, and probed for TOPBP1 occupancy by Western blotting. As shown in Figure 1C, both wild type and S1131A TOPBP1 could bind to DSBs. We conclude S1131 is dispensable for recruitment of TOPBP1 to DSBs, and likely functions after recruitment to stimulate DNA-bound ATR kinase.

What is the role of S1131 in ATR activation? Previous work has shown that phosphorylation of S1131 (to produce P-S1131) makes binding of TOPBP1 to ATR-ATRIP more efficient (Yoo et al, 2007). More recently, it has been shown for human TOPBP1 that P-S1138, the analogous residue, is required for TOPBP1’s ability to phase separate (Frattini et al, 2021). This, in turn, promotes robust ATR signaling. Importantly, although the human AAD contains S1138, the AAD by itself cannot undergo phase transition as additional sequences are required (Frattini et al, 2021). The AAD by itself can, however, activate ATR when added as a GST fusion protein to XEE (Kumagai et al, 2006). Given these circumstances, we reasoned that if S1131 functions exclusively in the context of condensate formation then it should be irrelevant to the ability of the isolated AAD, which cannot form a condensate, to activate ATR. To put this notion to the test, we prepared GST-AAD fusion proteins of three types: wild type, S1131A, and S1131D. Previous work has shown that the S1131D substitution, which mimics P-S1131, produces a dominant, gain-of-function TOPBP1 that binds ATR-ATRIP more efficiently and activates ATR more readily (Yoo et al, 2007). This previous work did not, however, test the S1131D mutation in the context of the isolated AAD. The three different GST-AAD proteins were added to XEE at the indicated concentrations and ATR signaling was assessed by anti P-CHK1 Western blotting (Figure 1D). The S1131A mutant produced less P-CHK1 than did the wild type version when both were present at 4 uM (compare lanes 2 and 4 in Figure 1D). This shows that S1131 is important for the ability of the isolated AAD to activate ATR, and thus that S1131 can function independently of condensate formation to promote ATR signaling. Interestingly, we observed the opposite effect for the S1131D mutation which hyper-activated ATR (compare lanes 2 and 5 in Figure 1D). This is consistent with previous data showing that S1131D in the context of full-length TOPBP1 promotes ectopic activation of ATR (Yoo et al, 2007), and our data extend these previous data by showing that S1131D can exert this effect in the context of the AAD alone. Based on the data in Figure 1, we conclude that the ability of S1131 to promote ATR activation is an intrinsic property of the AAD, and thus can be disconnected from condensate formation. This is not to say that it is not involved in condensate formation, but it is clearly doing something separate from phase transition.

### 3.2 S1131 promotes TOPBP1 multimerization

In recent work we used a synthetic form of TOPBP1, TOPBP1 Jr, to show that just three functional domains are needed for regulated activation of ATR by TOPBP1 (Ruis et al, 2021). The BRCT0-2 domains mediate recruitment and are dispensable thereafter while the AAD interfaces with ATR-ATRIP and the BRCT7&8 domains promote multimerization in an unknown manner. To determine if S1131 is possibly connected to TOPBP1 multimerization we first asked if the BRCT7&8 domains of TOPBP1 could interact with the AAD. For this, a standard GST pull-down assay and the GST fusion proteins we used are shown in Figure 2A. These proteins were used in binding assays with myc-tagged BRCT7&8, produced by IVTT. As shown in Figure 2B, myc-BRCT7&8 did not bind to GST alone, but did bind to GST-AAD, GST-BRCT7&8, and a fragment containing both the AAD and BRCT7&8. We next examined how the same set of GST proteins would bind to myc-tagged AAD alone, and again detectable binding was observed for all of the GST fusions but not GST alone (Figure 2C). In this experiment, however, the binding efficiencies differed, with the more efficient binding observed for GST-BRCT7&8, less efficient binding to GST-AAD-BRCT7&8, and relatively weak binding to GST-AAD. These data show that the AAD domain interacts with itself weakly, and that it binds BRCT7&8 more robustly (Figure 2D). In addition, the data show that BRCT7&8 can self-associate. We note that others have previously shown that the AAD can interact with itself (Thada et al, 2019).

**Figure 2.**
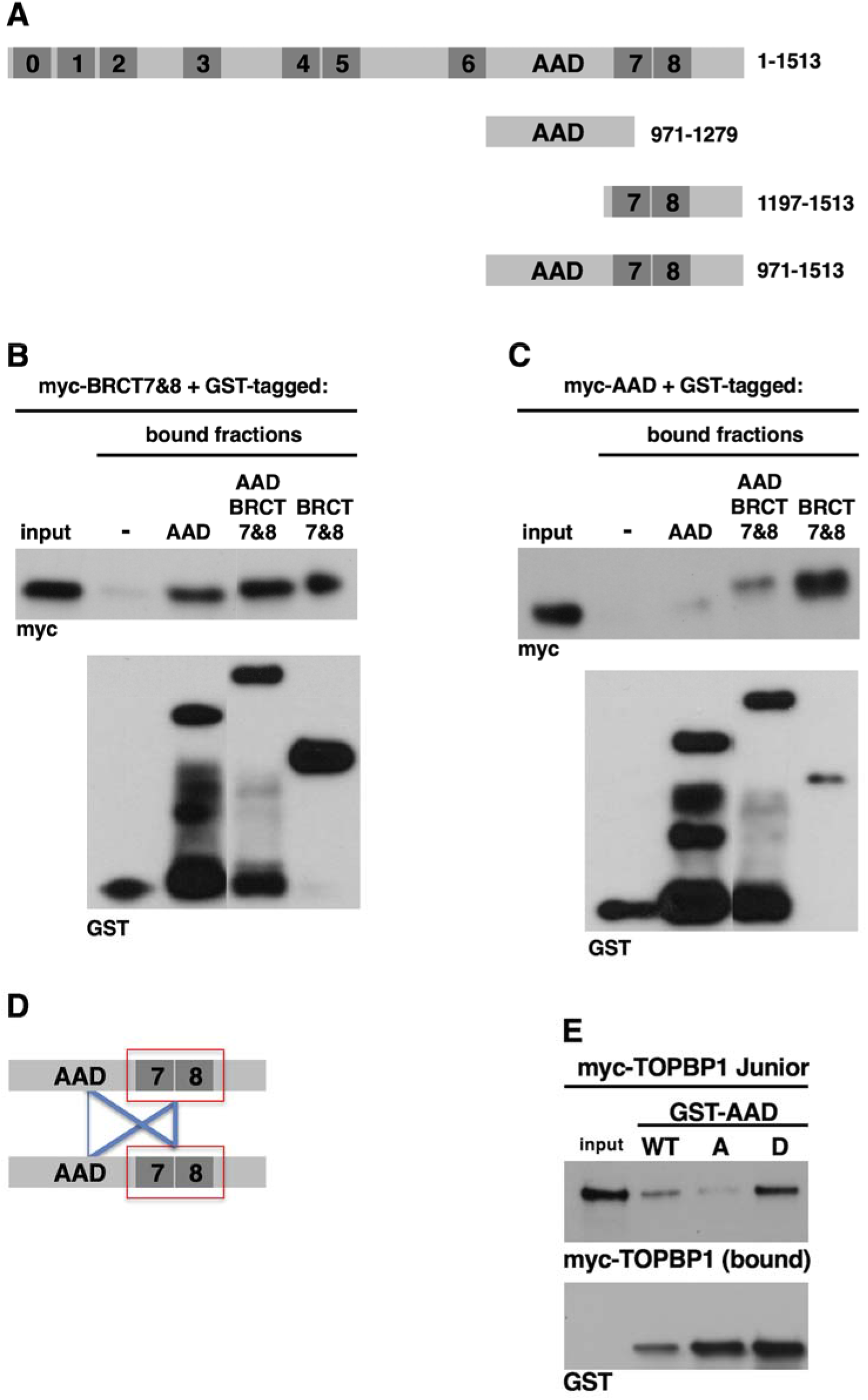
S1131 is important for multimerization of the AAD. **A.** Schematic representation of the proteins used for protein binding assays. **B.** The indicated GST proteins, expressed and purified from *E. coli*, were incubated in IVTT reactions expressing myc-tagged BRCT7&8. After incubation the samples were diluted in Binding Buffer and incubated with glutathione-sepharose beads. The beads were then washed, and the samples were eluted and probed by Western blotting for the indicated proteins. A sample of the total binding reaction, taken prior to dilution, was also probed for myc (lane “input”). The experiment shown is representative of two independently performed biological replicates. **C.** Same as (B) except myc-AAD was used. The experiment shown is representative of two independently performed biological replicates. **D.** Schematic showing how the AAD-BRCT7&8 region interacts with itself. **E.** Same as (B) except myc-TOPBP1 Jr was used with the indicated GST fusion proteins. The experiment shown is representative of two independently performed biological replicates.

To pursue these observations, we next asked how S1131 would impact interaction of the ADD with the TOPBP1 Jr. We chose TOPBP1 Jr for this analysis because, unlike the other constructs, TOPBP1 Jr has been shown to activate ATR in the context of DSBs (Ruis et al, 2021). We used the same GST proteins for this experiment as were used in Figure 1D. As shown in Figure 2E, myc-tagged TOPBP1 Jr could bind to the wild type AAD, and binding was somewhat reduced with the S1131A mutant. Interestingly, binding was increased, relative to wild type, with the S1131D mutant. Thus the same phospho-mimic mutation that increases ATR activation by the AAD also increases multimerization. We conclude that promoting multimerization is a previously undisclosed function of S1131.

### 3.3 T1098 within the AAD is phosphorylated in XEE and required for ATR activation

During the course of our studies of TOPBP1’s AAD, we noticed that, after incubation in XEE, that GST-AAD protein migrated with reduced mobility on SDS-PAGE. This can be seen in Figure 3A, where GST-AAD that had been incubated in XEE and then recovered back out of the extract using glutathione sepharose beads was run side-by-side with the input material. This reduced mobility suggests that post-translational modifications (PTMs) had been placed on GST-AAD during incubation in XEE, and thus to gain a sense of what these PTMs are we excised the band in Figure 3A, lane 3, and submitted it to analysis by mass spectrometry. This analysis revealed several peptides containing phosphorylated serine or threonine residues (Figure 3B). Figure 3C shows the sequence of the AAD from *Xenopus* TOPBP1 and these candidate phospho-acceptor sites are highlighted in red. The underlined portion shows the region in the frog protein that corresponds to the minimally defined AAD from human TOPBP1 (Thada et al, 2019), and thus two of the candidate phospho-acceptor sites (S1095 and T1098) fall within this region. We note that S1131 is only phosphorylated during a DNA damage response (Yoo et al, 2007), and thus we did not expect to recover P-S1131 signals in this experiment.

**Figure 3.**
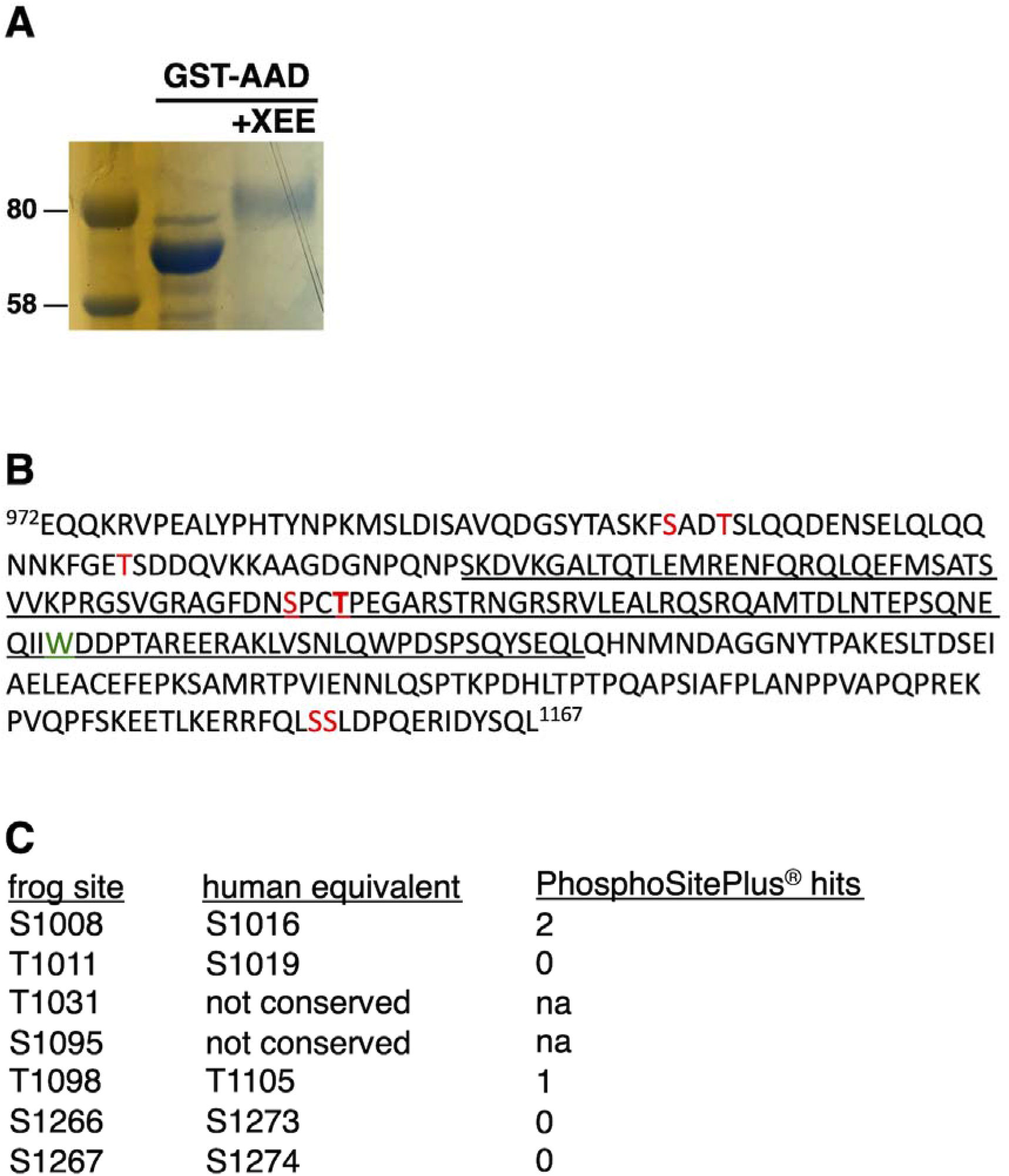
Mass spectrometry reveals multiple phospho-acceptor sies within the AAD region. **A.** A Coomassie blue stained SDS-PAGE gel showing altered mobility of GST-AAD after incubation in XEE. **B.** The amino acid sequence of the AAD region used for mass spectrometry analysis. See main text for details. **C.** Summary of conservation and PhosphoSitePlus® hits for the phospho-acceptor sites identified by mass spectrometry.

To pursue these observations, we next asked two questions – are the candidate phospho-acceptor sites identified by the mass spectrometry analysis conserved in human TOPBP1 and, if so, have any of those sites been curated on the website PhosphoSitePlus® (www.phosphosite.org), which catalogs phospho-sites for all human proteins? The answers are displayed in Figure 3D, where seven of the nine sites identified by mass spectrometry are conserved in human TOPBP1 and two of these, S1008 and T1098, are listed on PhosphoSitePlus®. Because T1098 is conserved in humans, listed on PhosphoSitePlus®, and falls within the minimally defined AAD, we focused on it first. The MS/MS spectrum for the peptide harboring pT1098 is displayed in Figure 4A. To determine if this residue is important for ATR activation, we produced a T1098A point mutant in full-length TOPBP1 and we tested this mutant for ATR activation, exactly as we did for S1131 in Figure 1B. As shown in Figure 4B, the T1098A mutant fails to activate ATR. This result shows that S1131 is not the only critical phospho-acceptor site within the AAD, as T1098 is also important.

**Figure 4.**
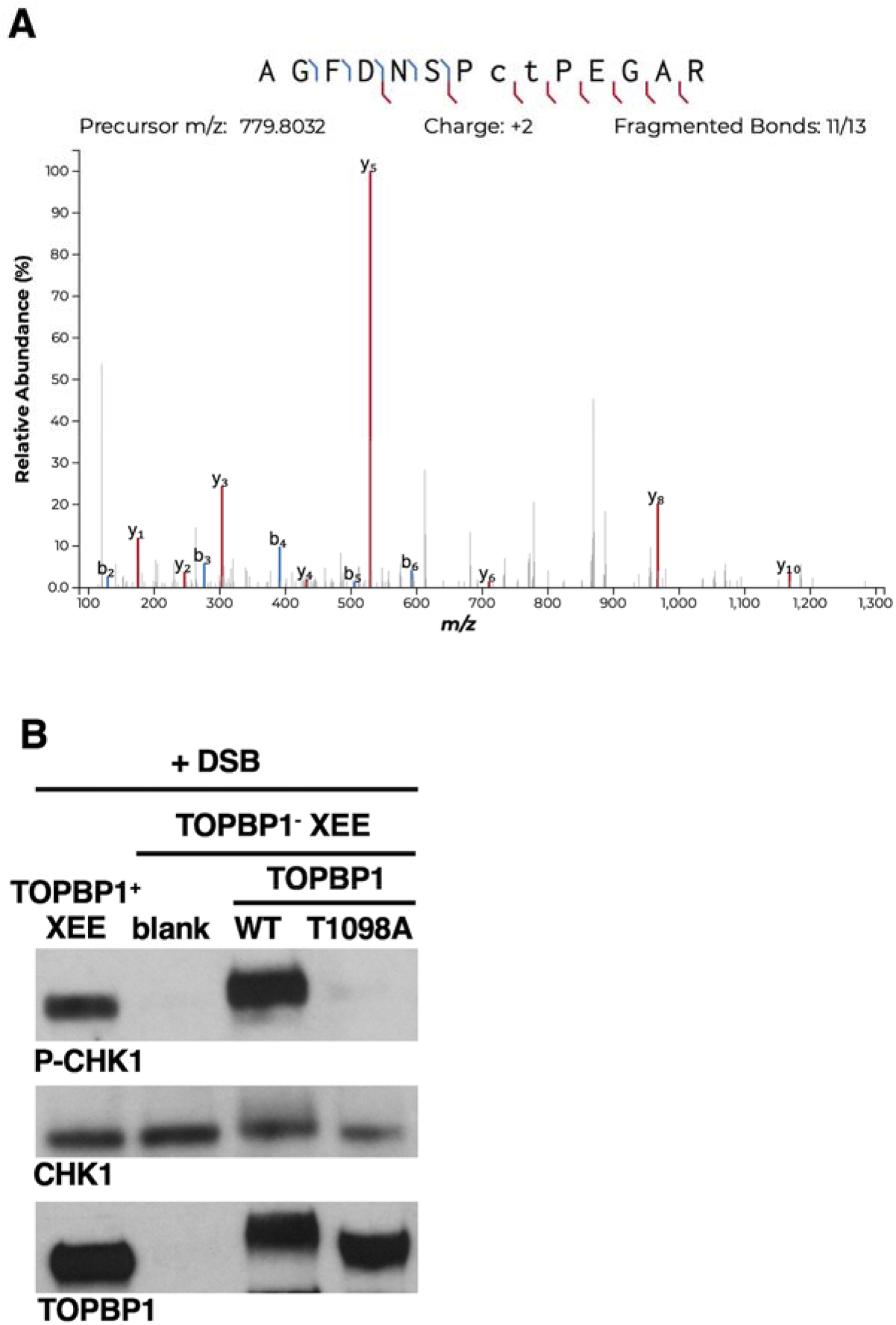
T1098 is required for ATR activation by TOPBP1. **A.** MS/MS spectra for the peptide harboring T1098. **B.** Same as Figure 1B except the T1098A mutant was used. The experiment shown is representative of two independently performed biological replicates.

### 3.4 A casein kinase 2 inhibitor blocks both DSB- and ADD-mediated ATR activation

One feature of the AAD region of TOPBP1 is that it contains an unusually high number of consensus sequences for phosphorylation by casein kinase 2. The consensus motif for casein kinase 2 is S/T-X-X-D/E (St-Denis et al, 2015), and this motif is found nine times in the AAD region (Figure 5A), and one of these (S1266) was identified in our mass spectrometry analysis and is conserved in human TOPBP1. This prompted us to ask if casein kinase 2 activity is required for the AAD to function. For this, we asked if DSBs could still activate ATR in XEEs treated with the casein kinase 2 inhibitor (CK2i) CX-4945. We found that CX-4945 could block ATR activation (Figure 5B). We have previously shown that CX-4945 does not impair the recruitment of TOPBP1 to DSBs (Montales et al, 2022) and thus we considered the possibility that the AAD requires casein kinase 2 activity in order to function. To test this idea we added GST-AAD to XEE and observed that CX-4945 could still inhibit ATR activation. Based on this experiment, we conclude that the AAD requires, either directly or indirectly, casein kinase 2 activity in order to activate ATR.

**Figure 5.**
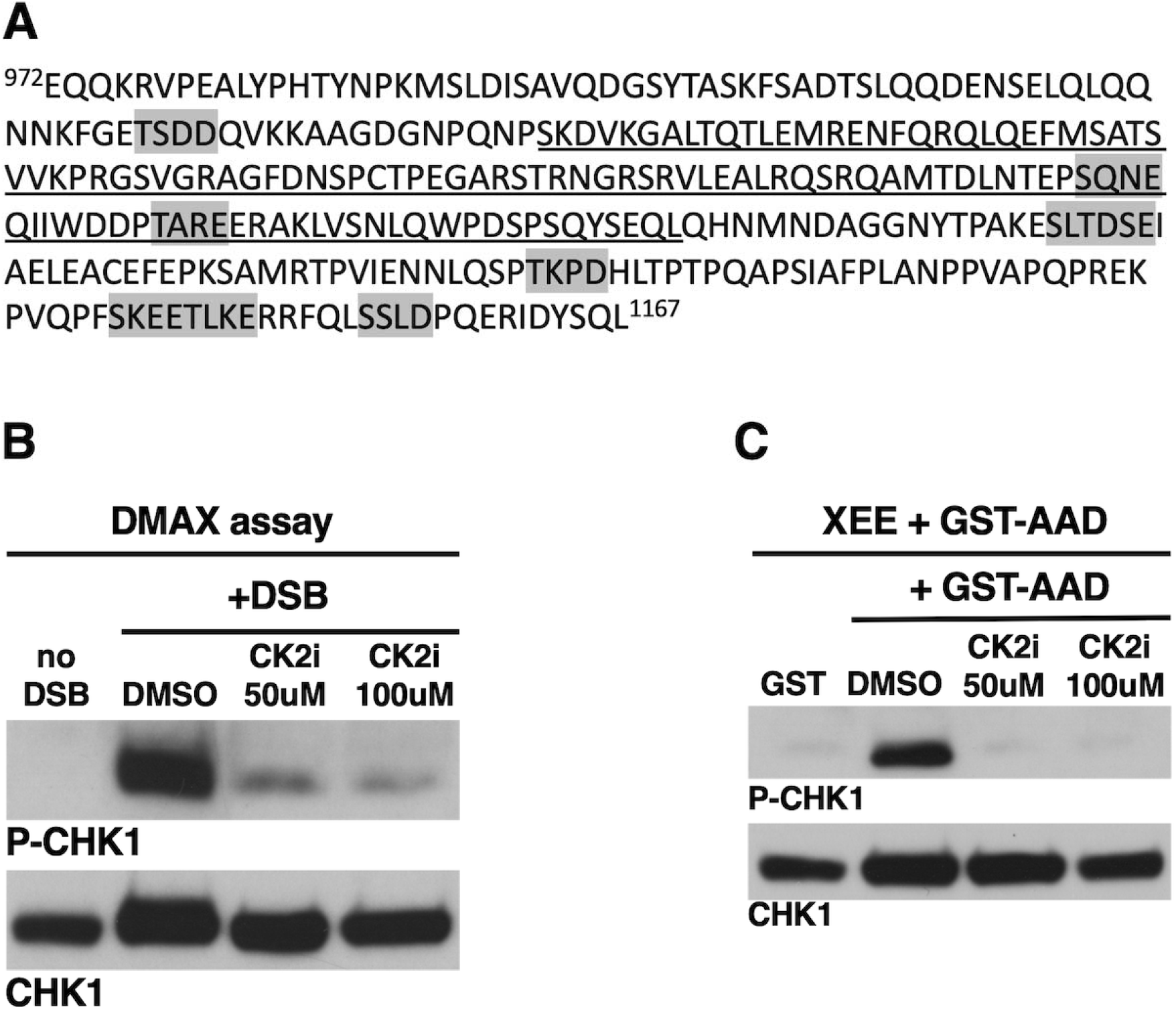
Delineation of the NBS1 binding region within TOPBP1’s BRCT1 domain. **A.** Same as Figure 3C except the position of consensus casein kinase 2 phosphorylation sites is highlighted with grey shading. **B-C.** XEEs were treated as indicated with either DMSO or CK2i and either DSBs (B.) or the indicated GST proteins (C.) were added. After incubation the samples were probed by Western blotting for P-CHK1 and CHK1. The experiments shown are representative of two independently performed biological replicates.

In this paper we present data showing that phosphorylation of TOPBP1’s AAD is crucial for its ability to activate ATR. Previous work had identified S1131 as a regulated phospho-acceptor site necessary for ATR activation by DSBs, but not stalled replication forks. This previous work showed that S1131 is phosphorylated by ATM, and that the manifestation of this phosphorylation was increased affinity for ATR-ATRIP (Yoo et al, 2007). Here, we extend these observations by showing that S1131 is important for multimerization of the AAD (Figure 3E). We also report that an additional residue within the AAD, T1098, is also phosphorylated, but this is constitutive phosphorylation as it occurs in the absence of a DDR response. Like S1131, phosphorylation of T1098 is required for ATR activation. Summarizing these findings, we see that under resting conditions, TOPBP1 is primed for action via constitutive phosphorylation on T1098; however in its monomeric form, it is unable to activate ATR (Figure 7). ATM-directed phosphorylation of S1131 occurs in response to DSBs, and now TOPBP1 can assume the multimeric form that is required for ATR activation (Figure 7). It is likely that additional sites within the AAD are also phosphorylated, as this region contains multiple casein kinase 2 consensus motifs and we have shown here that casein kinase 2 activity is required for AAD function (Figure 6). In conclusion, our data show that the AAD undergoes multi-site phosphorylation, under both constitutive and regulated conditions, and that at least some of these phosphorylations convert TOPBP1 from an inactive to an active form via multimerization.

**Figure 6.**
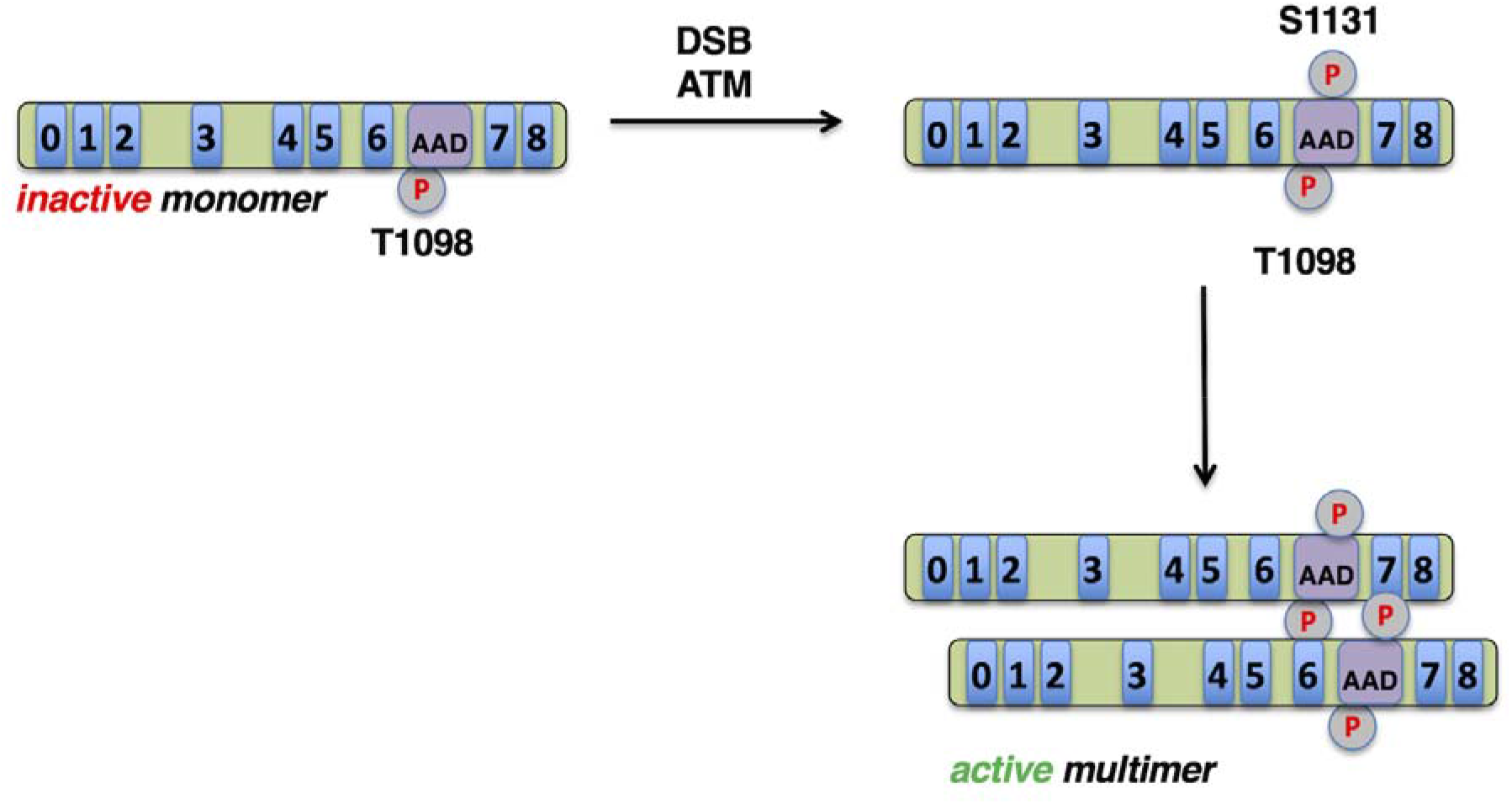
Multi-site phosphorylation of the TOPBP1 AAD drives ATR activation through a multimerization mechanism. Please see text for details.

## Author contribution

W.M.M. conceptualized the project, acquired the funding, and wrote the manuscript. H.S., K.R., and K.M. performed the experiments. J.A.W. and Y.J. performed the mass spectrometry experiments.

## Acknowledgments

This work was supported by NIH grant R01GM12287 (W.M.M.) and R01GM089778 (J.A.W.). We thank Shan Yan and Hovik Gasparyan for the production of plasmid constructs used in this study.

